# A simple linearization method unveils hidden enzymatic assay interferences

**DOI:** 10.1101/528596

**Authors:** Maria Filipa Pinto, Jorge Ripoll-Rozada, Helena Ramos, Emma E. Watson, Charlotte Franck, Richard J. Payne, Lucília Saraiva, Pedro José Barbosa Pereira, Annalisa Pastore, Fernando Rocha, Pedro M. Martins

**Affiliations:** Instituto de Ciências Biomédicas Abel Salazar (ICBAS), Universidade do Porto, Porto, Portugal; Laboratório de Engenharia de Processos, Ambiente, Biotecnologia e Energia (LEPABE), Faculdade de Engenharia da Universidade do Porto (FEUP), Porto, Portugal; IBMC - Instituto de Biologia Molecular e Celular, Universidade do Porto, Porto, Portugal; Instituto de Investigação e Inovação em Saúde, Universidade do Porto, Porto, Portugal; Laboratório Associado para a Química Verde (LAQV), Rede de Química e Tecnologia (REQUIMTE), Faculdade de Farmácia da Universidade do Porto (FFUP), Porto, Portugal; School of Chemistry, The University of Sydney, Sydney, New South Wales 2006, Australia; Maurice Wohl Clinical Neuroscience Institute, King’s College London, London, England; Scuola Normale di Pisa, Pisa, Italy

## Abstract

Enzymes are among the most important drug targets in the pharmaceutical industry. The bioassays used to screen enzyme modulators can be affected by unaccounted interferences such as time-dependent inactivation and inhibition effects. Using procaspase-3, caspase-3, and α-thrombin as model enzymes, we show that some of these effects are not eliminated by merely ignoring the reaction phases that follow initial-rate measurements. We thus propose a linearization method (LM) for detecting spurious changes of enzymatic activity based on the representation of progress curves in modified coordinates. This method is highly sensitive to signal readout distortions, thereby allowing rigorous selection of valid kinetic data. The method allows the detection of assay interferences even when their occurrence is not suspected *a priori*. By knowing the assets and liabilities of the bioassay, enzymology results can be reported with enhanced reproducibility and accuracy. Critical analysis of full progress curves is expected to help discriminating experimental artifacts from true mechanisms of enzymatic inhibition.

## Introduction

Typically, more than one-third of the discrete drug targets in the portfolio of pharmaceutical companies consists of enzymes [1], with phosphate-transferring enzymes, or kinases, being the largest category of potentially novel drug targets [2]. Drug screening is usually based on enzymatic assays that aim at identifying compounds that inhibit, enhance or modulate enzyme activity. However, the output of these assays strongly depends on the experimental conditions and on several different parameters that are often difficult to master completely. In high-throughput screening (HTS) of enzyme modulators, primary assays employing light-based detection methods are escorted by orthogonal assays using different output reporters in order to identify false positives and fluorescence/luminescence artifacts [3, 4]. Other possible interferences can be specific of a given system, such as the occurrence of enzyme inactivation and competitive product inhibition, or unspecific, as in the cases of random experimental errors and of changes in experimental parameters during the reaction (Figure 1). This uncertainty dramatically calls for new and more sensitive approaches to allow fast and reliable detection of these interferences.

**Figure 1.**
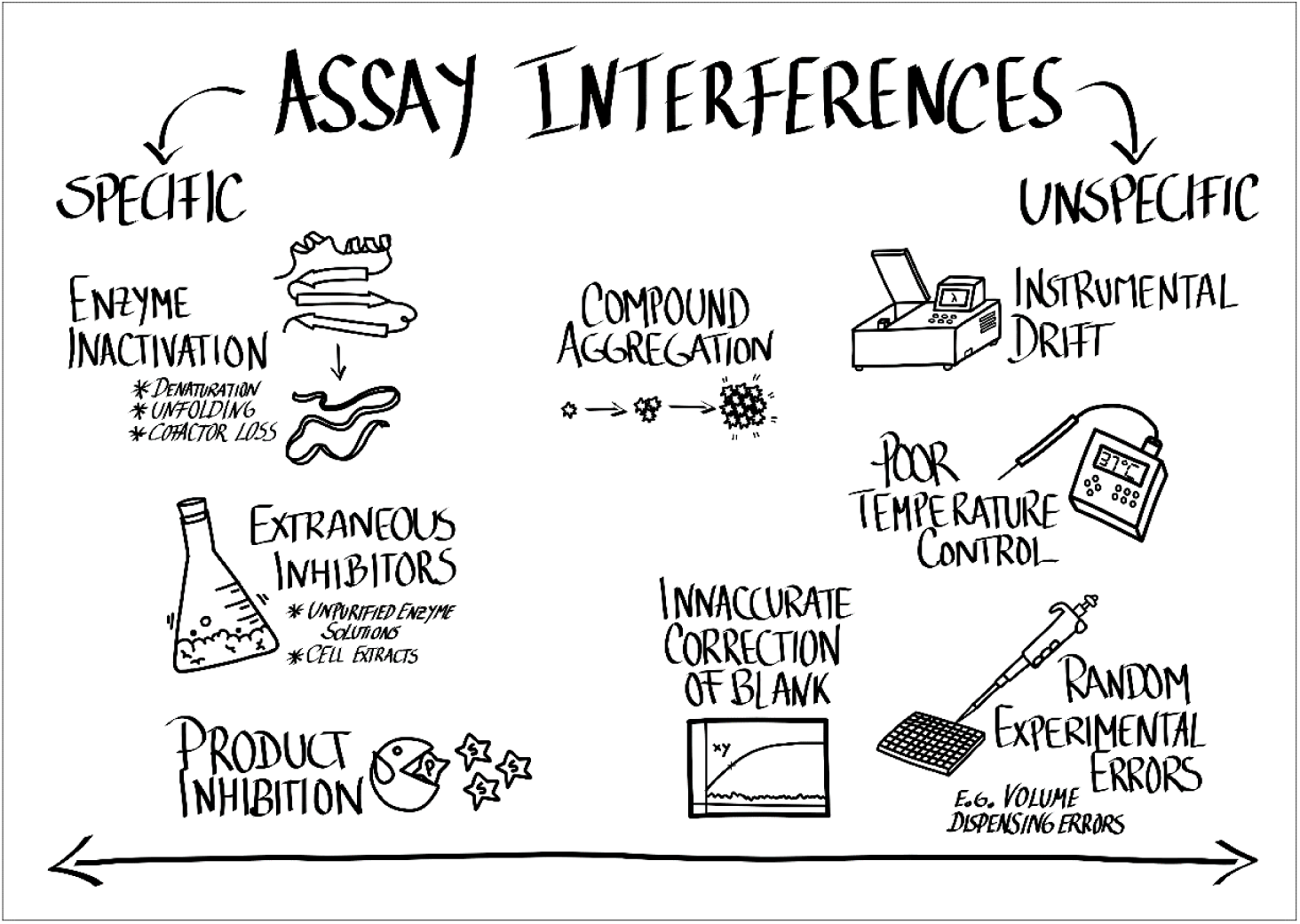
Specific and unspecific interferences on enzymatic assays. Spurious effects that are too small to be readily observable can produce important errors of interpretation of kinetic results. Specific of a given system, enzyme inactivation can be prevented by the addition of protein stabilizers or by increasing the concentration of enzyme [5]. Extraneous inhibitors present in unpurified enzyme solutions or in cell extracts systematically affect the quality of kinetic measurements [6]. Product inhibition can usually be ignored in initial-rate measurements but is highly misleading in time-course studies [7]. Compound aggregation can cause enzyme sequestration on the surface of the aggregate particles and is one of the main reasons for promiscuous enzyme inhibition [3]. Instrumental drift, poor temperature control, inaccurate correction of the sample blank, and random experimental errors in e.g. volume dispensing operations are typical examples of unspecific interferences [8, 9]. Adequate buffering of the reaction mixture is important to prevent changes of enzyme activity provoked by drifts in pH and ionic strength [10].

In the present contribution, we propose a touchstone criterion for the detection of assay interferences based on the graphical representation of reaction coordinates in a linearized scale. We applied our method to enzymatic reactions catalyzed by procaspase-3, caspase-3 and α-thrombin. Caspases are a family of cysteine-dependent aspartate-specific proteases (MEROPS family C14; [11]) synthesized as zymogens and converted into their more active forms upon proteolytic cleavage [12]. Both caspase-3 and its precursor procaspase-3 undergo progressive inactivation during *in vitro* enzymatic assays. Progress curves of procaspase-3- and caspase-3-catalyzed reactions are analyzed to identify enzyme inactivation and characterize its relative importance. Alpha-thrombin is a (chymo)trypsin-like serine proteinase (MEROPS family S01; [11]) and a main effector in the coagulation cascade. Similar to caspase-3, its zymogen (prothrombin) is cleaved to generate the active form of the enzyme. Thrombin generation is tightly regulated to allow blood clot formation after an injury [13]. A variety of thrombin-targeting inhibitors is produced by blood-feeding organisms [14-17]. The outcome of the new test in the presence of enzyme inhibition is demonstrated for α-thrombin-catalyzed reactions inhibited by a synthetic variant of an anticoagulant produced by *Dermacentor andersoni* [18, 19]. Along with the inactivation and inhibition studies, we discuss the detection of unspecific interferences arising from changes in the reaction conditions.

## Experimental procedures

### Procaspase-3 production in yeast cell extracts

Procaspase-3 was produced as previously described [20, 21]. Briefly, cultures of *Saccharomyces cerevisiae* transformed with the expression vector pGALL-(*LEU2*) encoding human procaspase-3 were diluted to 0.05 optical density at 600 nm (OD_600_) in 2% (w/v) galactose selective medium and grown at 30°C under continuous shaking until an OD_600_ range of 0.35-0.40. Cells were collected by centrifugation and frozen at −80°C. For protein extraction, cell pellets were thawed, treated with *Arthrobacter luteus* lyticase (Sigma-Aldrich, Sintra, Portugal), and the cells were lysed using CelLytic™ Y Cell Lysis Reagent (Sigma-Aldrich, Sintra, Portugal) in the presence of 1 mM phenylmethylsulfonyl fluoride, 1 mM Dithiothreitol (DTT), 1 mM ethylenediaminetetraacetic acid (EDTA), 0.01% (v/v) Triton X-100. Total protein concentration of the extracts was determined using the Pierce™ Coomassie Protein Assay Kit (ThermoFisher Scientific, Porto Salvo, Portugal).

### Enzymatic assays for procaspase-3 and caspase-3

The activity of recombinant human procaspase-3 and of recombinant human purified caspase-3 (Enzo Life Sciences, Lisboa, Portugal) was followed by monitoring the conversion of the fluorogenic substrate Acetyl-Asp-Glu-Val-Asp-7-amido-4-methylcoumarin (Ac-DEVD-AMC, Sigma-Aldrich, Sintra, Portugal) to 7-amino-4-methylcoumarin (AMC) at 37°C. Procaspase-3 (0.123 mg/mL protein extract) and caspase-3 (1.0 U) were assayed in 96-well microplates (Nunc™ MicroWell™ 96-Well, Thermo Scientific™, ThermoFisher Scientific, Porto Salvo, Portugal) using 100 μL of 20 mM HEPES pH 7.4, 100 mM NaCl, 10% (w/v) sucrose, 0.1% (w/v) CHAPS, 10 mM DTT, 1 mM EDTA per well [22]. A range of substrate concentrations of 0-300 μM Ac-DEVD-AMC for procaspase-3, and 0-50 μM Ac-DEVD-AMC for caspase-3 were tested. The reactions were started by addition of protein, and fluorescence was monitored at 460 nm (390 nm excitation) using a HIDEX CHAMELEON V plate reader (Turku, Finland). To avoid evaporation, the reaction mixture in each well was overlaid with liquid paraffin (100 μL). All solutions were equilibrated to 37°C before use. Assays were performed in triplicate or quadruplicate for procaspase-3, and in duplicate for purified caspase-3. The calibration curve was built using solutions with known concentration of the free fluorescent product AMC (Sigma-Aldrich, Sintra, Portugal).

### Enzymatic assay for α-thrombin

The enzymatic activity of human α-thrombin (Haematologic Technologies, Essex, USA) was assessed by following its amidolytic activity toward the chromogenic substrate Tos-Gly-Pro-Arg-p-nitroanilide (Chromozym TH, Roche, Amadora, Portugal) at 37°C in the presence of 0.40 nM of a synthetic variant of an anticoagulant produced by *D. andersoni* [19]. The assays were performed at least in duplicate, as previously described, except for the concentration of α-thrombin (0.15 nM) [23].

## The linearization method

Deviations from the normal progress of enzyme-catalyzed reactions should, in principle, alter the build-up profile of product concentration (*P*) vs. time (*t*) from the theoretical curve expected by the integrated form of the Michaelis-Menten (MM) equation [24-26]:

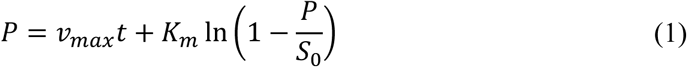

where *S*_0_ is the initial substrate concentration and *K*_*M*_ and *v*_*max*_ are the MM constants. In practice, however, Eq. 1 is not used to detect assay interferences since no evident changes in the shape of the progress curves are induced by enzyme inactivation, product inhibition, and quasi-equilibrium mechanisms of competitive inhibition, uncompetitive inhibition, etc. [27]. In the specific case of enzyme inactivation interferences, their occurrence can be detected by the Selwyn test applied to progress curves measured at different enzyme concentrations (*E*_0_) and constant *S*_0_ [28]. Yet, the Selwyn test might not detect cases of incomplete enzyme inactivation, besides requiring the realization of additional experiments. Eq. 1 produces estimates of kinetic parameters poorer than the classic MM equation fitted to initial reaction rate measurements because any external interference may severely accumulate over the full time-course [25, 26]. In addition, the application of Eq. 1 is limited to a range of conditions more restricted than that of *E*_0_ ≪ *S*_0_ + *K*_*m*_ required to validate the steady-state assumption [29, 30]: Eq. 1 fails to account for the fast transient phase that precedes the steady-state phase according to the closed-form solution of the Briggs-Haldane reaction mechanism (Figure 2a). Additionally, the effect of *E*_0_ can be masked by apparent values of the Michaelis constant 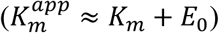, especially for *E*_0_ ≥ *K*_*m*_ [31].

**Figure 2.**
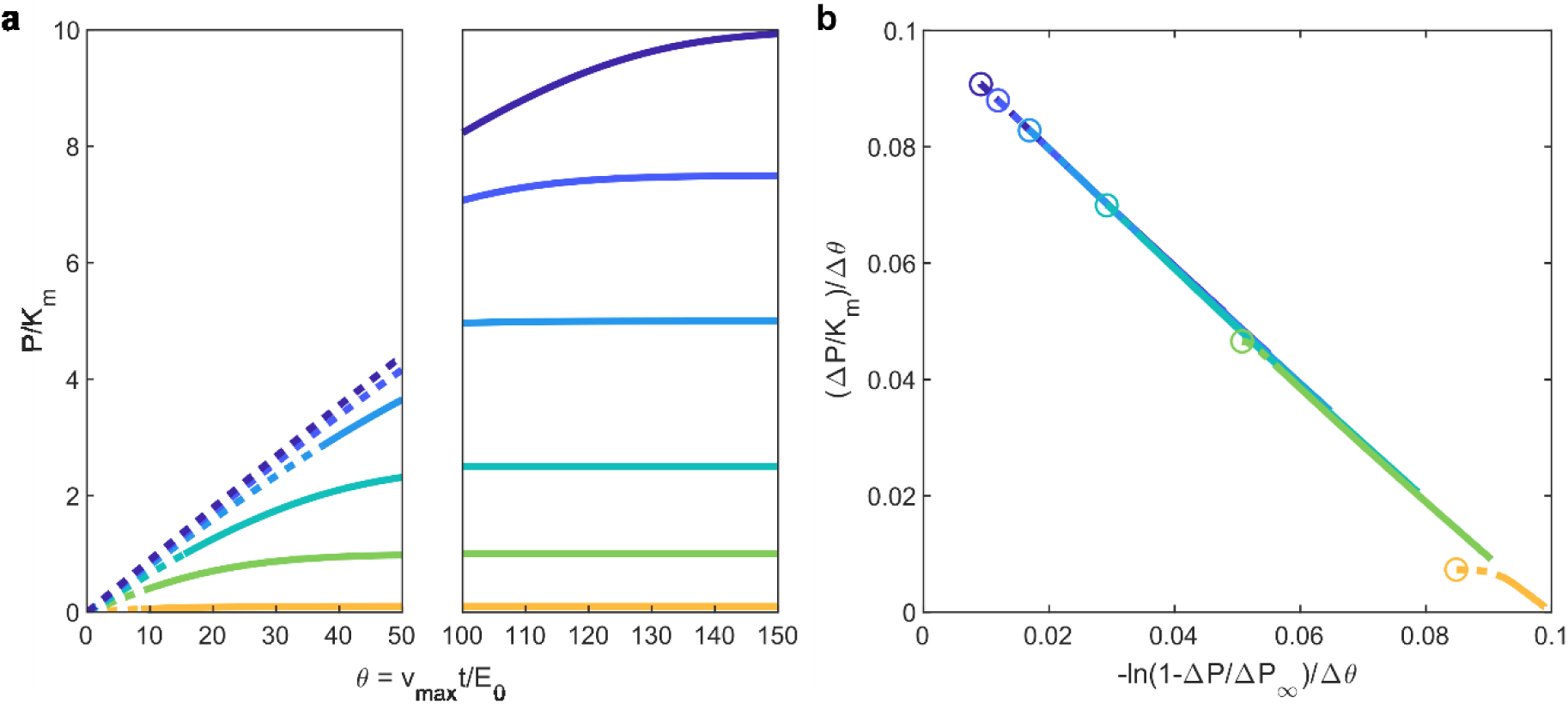
Theoretical progress curves expected for unbiased Briggs-Haldane reaction mechanisms. (a) Evolution of product concentration over time represented in dimensionless units of product concentration (*p* = *P*/*K*_*m*_) and time (*θ* = *v*_*max*_ *t*/*E*_0_) for different substrate concentrations. Different colors correspond to values of *s*_0_ = *S*_0_/*K*_*m*_ of (from top to bottom) 10, 7.5, 5, 2.5, 1, 0.1 (additional simulation parameters given in Table A1, Appendix). The broken *x* axis is used to emphasize the initial periods of constant velocity (dashed lines). (b) The same curves are represented in linearized coordinates according to Eq. 2; round markers indicate the steady-state instant *t*\*. The absence of assay interferences is evidenced by negatively-sloped, superimposing straight lines.

We propose a new linearization method (LM) for the detection of assay interferences based on the following modified version of the integrated MM equation:

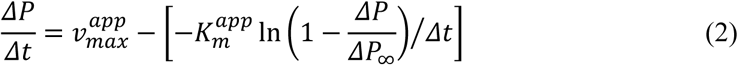

The main differences of this formalism relatively to Eq. 1 are: (a) the use of apparent kinetic constants 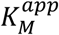 and 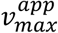, (b) the use of partial time intervals (Δ*t* = *t* − *t*_*i*_) and of the corresponding increment of product concentration (Δ*P* = *P* − *P*_*i*_), (c) the initial condition (subscript *i*) is now any point of the reaction subsequent to the pre-steady-state phase (*t*_*i*_ > *t*\*), (d) the final concentration of product is given by the measured value (*P*_∞_) and not by the expected value (*S*_0_) [26], and (e) Eq. 2 is presented in a linearized form of the Walker and Schmidt type [26] (Figure 2b). Features (a) to (c) are meant to expand the validity of time-course analysis to the same range of conditions of the steady-state assumption (*E*_0_ ≪ *S*_0_ + *K*_*m*_) [31, 32]. Feature (d) takes into account possible discrepancies between *P*_∞_ and *S*_0_ values resulting, for example, from complete enzyme inactivation or inaccurate pipetting. Feature (e) is implemented because the Walker and Schmidt linearization is highly sensitive to fluctuations in *P* readouts [33], whereas linearity is an easily implementable judgment criterion.

The linear variation of *ΔP*/*Δt* with −ln(1 − *ΔP*/*ΔP*_∞_)/*Δt* expected by Eq. 2 is a necessary but not sufficient condition to reject assay interferences. To pass this test, the straight lines obtained at different substrate concentrations should also superimpose each other (Figure 2b). Although parameter estimation is not the primary goal here, the agreement between the apparent constants and the values of *K*_*m*_ and *v*_*max*_ obtained by the standard initial-rate method further confirms that the assay is unbiased. Finally, since the LM equation applies to single active site, single substrate and irreversible steady-state reactions of the Briggs-Haldane type [34], this method might also be used to reveal the presence of a more complex enzymatic mechanism from the one assumed.

## Results and discussion

### Procaspase-3 inactivation - preliminary analysis

The hyperbolic reaction curves of Ac-DEVD-AMC cleavage by procaspase-3 (Figure 3a) are not suggestive of any evident loss of enzyme activity over time. The lack of well-defined slopes from which the initial rates (*v*_0_) can be accurately measured might only indicate that the substrate concentrations are still too low to achieve the saturating MM conditions [35, 36]. In fact, the plateau corresponding to *v*_*max*_ in the MM representation (Figure 3b) is barely noticeable in the studied range of substrate concentrations.

**Figure 3.**
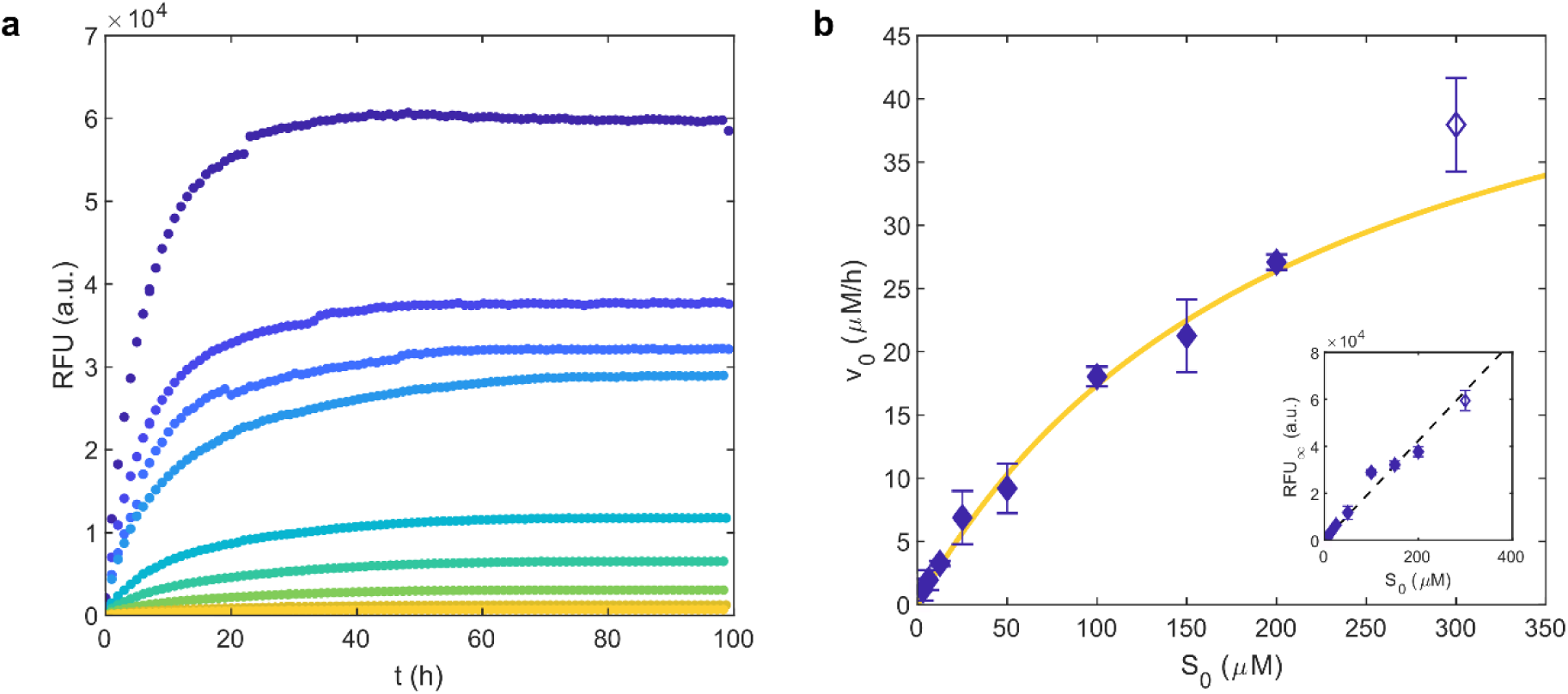
Loss of procaspase-3 activity is not self-evident in a first analysis of kinetic results. (a) Fluorescence (*RFU*) increase during the cleavage of Ac-DEVD-AMC by procaspase-3 for *S*_0_ values of (from top to bottom) 300, 200, 150, 100, 50, 25, 12.5, 6.25, and 3.125 μM. (b) Plot of the initial reaction rates (*v*_0_) as a function of substrate concentration. Symbols and error bars: means and standard deviations of *v*_0_ values calculated using the initial slopes obtained in (a) and a preliminary calibration curve relating the end-point fluorescence (*RFU*_∞_) and *S*_0_ (inset). Solid line: fit of selected experimental data (closed symbols) to the MM equation. Since the results are affected by severe enzyme inactivation, the fitted values of *K*_*m*_ = 217 ± 59 μM and *v*_*max*_ = 55 ± 9 μM are merely indicative. Inset: linear fit (dashed line) to selected (closed symbols) *RFU*_∞_ vs. *S*_0_ data.

The obtained value of *K*_*m*_ = 217 ± 59 μM is 62-fold higher than that previously reported using the uncleavable mutant procaspase-3(D_3_A), which has three processing sites removed, and the substrate Ac-DEVD-AFC, which has a different fluorescent reporter (AFC) than Ac-DEVD-AMC [37]. Differences between the observed and the literature values of *K*_*m*_ could therefore be ascribed to the distinct nature of each enzymatic assay. Yet, the calibration curve represented in Figure 3b (inset) provides important clues as to the possible existence of experimental artifacts. This is a conditional calibration curve since the final fluorescence value (*RFU*_∞_) is assumed to result from a complete catalytic reaction in which the final product concentration *P*_∞_ is equivalent to *S*_0_. To check the validity of this hypothesis, reference fluorophore solutions were used to calibrate the equipment according to the standard protocol (Figure 4a); for the same concentrations of fluorophore and substrate, the fluorescence intensity of the calibration solutions (*RFU*_cal_) clearly surpasses the *RFU*_∞_ signal, thus suggesting partial conversion of substrate into products during the reactions (Figure 3a). This finding invalidates the preliminary calibration curve and the kinetic analysis in Figure 3b since the condition of complete chemical reaction is not observed. Whether or not the reactions were really unfinished and what were the mechanisms thereby involved cannot be ascertained by the calibration-curve test alone. Besides procaspase-3 inactivation, other interfering factors, such as fluorescence quenching phenomena, could have caused the observed differences between *RFU*_cal_ and *RFU*_∞_. The occurrence of progressive loss of enzyme activity was further confirmed by the Selwyn test (Figure 4b) and by the direct measurement of procaspase-3 activity for different periods of incubation in the reaction environment (Figure 4c). Relatively to these methods, the calibration-curve test (Figure 4a) has the advantage of requiring no other experiments than those already performed while estimating MM kinetic parameters. Furthermore, if enzyme inactivation is admitted as a first-order decay process, the following relationship can be used to quantitatively estimate the product of the decay rate constant *λ* by the time constant *τ* = *K*_*M*_/*v*_*max*_ (Appendix):

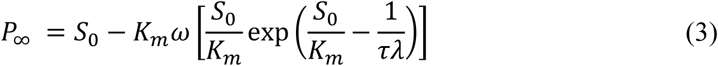

**Figure 4.**
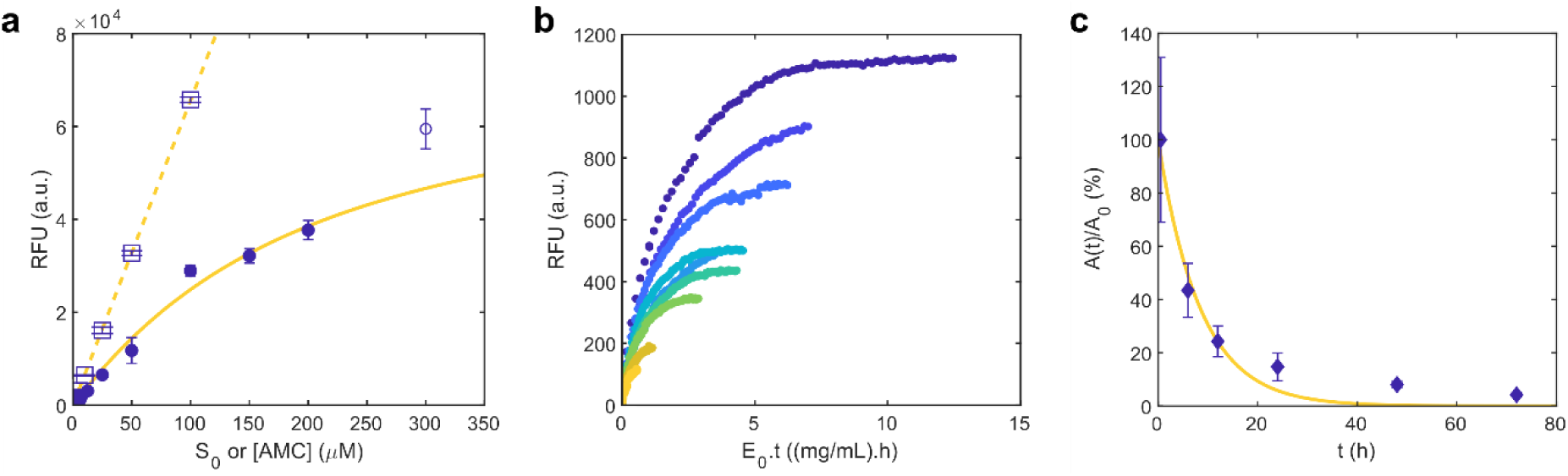
Procaspase-3 inactivation identified by different methods. (a) The calibration-curve test is proposed based on the differences between *expected* (*RFU*_cal_, squares) and *obtained* (*RFU*_∞_, circles) end-point signals. Symbols and error bars represent means and standard deviations. The values of *RFU*_cal_ are measured using solutions of known fluorophore concentrations ([AMC]). The test passes if the values of *RFU*_cal_ and *RFU*_∞_ superimpose. Solid line: the experimental data is fitted by Eq. 3 (fitting results: *K*_*m*_ = 161 ± 77 μM and *τλ* = 1.4 ± 0.3 h). The obtained end-point signals are the same used to build the preliminary calibration curve in the inset of Figure 3b. Dashed line: linear fit representing the true calibration curve. (b) The classic Selwyn test also suggests time-dependent loss of procaspase-3 activity since progress curves measured for (from top to bottom) 0.17, 0.13, 0.09, 0.06, 0.07, 0.06, 0.04, 0.02, 0.01 mg/mL procaspase-3 and constant *S*_0_ (3.125 μM) are not superimposable when represented in a modified *E*_0_*t* timescale [28]. (c) Symbols and error bars: means and standard deviations of the normalized enzymatic activity *A*(*t*)/*A*_0_ after different periods of incubation. Solid line: numerical fit to an exponential decay function (fitted result: *λ* = 0.12 ± 0.02 h^-1^).

For a given substrate concentration, the extent of the reaction is ultimately determined by the product *τλ* relating the rates of inactivation and of unimpaired reaction. It follows from the inverse dependence of *τλ* on *E*_0_ (via *τ* and *v*_*max*_) that the effects of inactivation can be counterbalanced by increasing the concentration of enzyme. Similar improvements can be achieved by decreasing the value of *λ* through the use of protein stabilizers. The fitted value of *τλ* = 1.4 ± 0.3 is clearly in the region above ∼0.1 for which complete enzyme inactivation can be attained before the conversion of the total available substrate (view Tables S2 and S3). Contrary to the estimation obtained by initial-rate analysis, the fitted value of *K*_*m*_ = 161 ± 77 μM is obtained taking into account the effect of enzyme inactivation. The quantitative information provided by the calibration-curve test is an advantage over the Selwyn test, whose underlying principle also requires additional progress curves to be measured with various enzyme concentrations and constant *S*_0_ values [28]. In the case described in Figure 4b, the non-superimposed Selwyn plots of *P* against *E*_0_*t* confirm the likely occurrence of procaspase-3 inactivation, as previously suggested by simple inspection of the calibration curves (Figure 4a). The last evidence supporting the verdict of both tests is obtained by directly measuring the enzymatic activity at the end of different periods of incubation in the reaction environment (Figure 4c). Procaspase-3 activity is confirmed to rapidly decrease with time according to the exponential decay trend expected for first-order processes. The reciprocal of the decay rate constant (1/*λ* = 8.3 h) can be interpreted as the period of time required for the catalytic activity to drop to ∼37% of its initial value.

### Loss of procaspase-3 activity identified by the linearization method

Complete catalytic reactions, promoted for example by high *E*_0_ values or by low *S*_0_ values, can pass the calibration-curve and the Selwyn tests even in presence of significant enzyme inactivation. This misjudgment may affect the overall quality of reported enzymology data, particularly when the initial reaction rate phase is itself difficult to define, as in conditions of *S*_0_ < *K*_*M*_ [35, 38]. Enhanced limits of detection to this and other interferences can be achieved by application of the new LM test brought forward by Eq. 2. In the commonest case of *E*_0_ ≪ *K*_*M*_, the intervals Δ*t* and Δ*P* can be considered right from the beginning of the measurements because the steady-state condition starts to be valid after the first milliseconds of the reaction [31, 39]. The non-conformity of the linearized curves of procaspase-3 (Figure 5) with the ideal behavior previously described in Figure 2b is evident: the linearized progress curves obtained for different values of *S*_0_ show positive slopes (rather than negative slopes) and do not superimpose.

**Figure 5.**
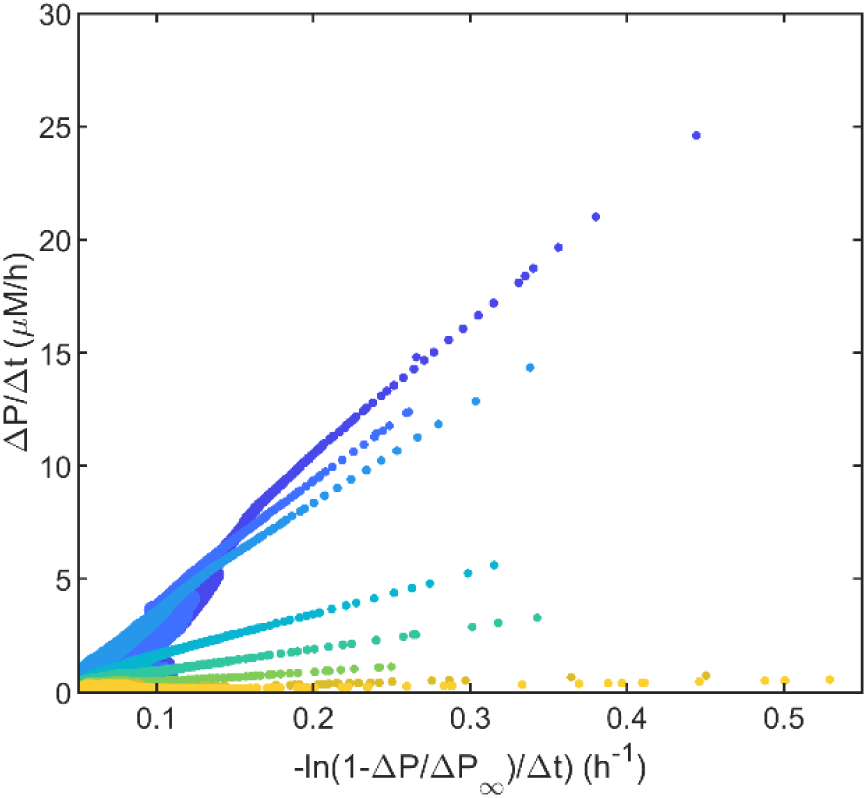
Using the LM test to detect procaspase-3 inactivation. The progress curves in Figure 3a measured for *S*_0_ values of 200, 150, 100, 50, 25, 12.5, 6.25, and 3.125 μM are now represented in the linearized Δ*P*/Δ*t* vs. −ln(1 − *ΔP*/*ΔP*_∞_)/*Δt* scale (color-coded as in Figure 3a). Symbol size increases with the time-course of the reaction. Non-superimposing, positively-sloped straight lines clearly indicate the occurrence of assay interferences, which, in the present case, are associated with procaspase-3 inactivation (Figure 4).

The calibration-curve test, the Selwyn test and the LM test are all capable of detecting the progressive inactivation of procaspase-3. For practical uses, the new methods here proposed are more easily applicable than having to prearrange and perform additional Selwyn test experiments. However, the question remains open as to which of the three methods is more sensitive to slight losses of enzymatic activity. In order to study milder decay processes than that observed for procaspase-3 in cell extracts, values of *τλ* < 1.4 were used in the numeric simulations described in detail in the Appendix (Tables S2-S4). For a reference value of *τλ* = 0.1 and assuming *S*_0_/*K*_*M*_ ratios between ∼1 and ∼5, both the calibration-curve test (Figure 6a) and the Selwyn test (Figure 6b) fail to reveal *S*_0_-dependent effects caused by inactivation. Conversely, these effects are visibly amplified in Figure 6c where the Δ*P*/Δ*t* vs. −ln(1 − *ΔP*/*ΔP*_∞_)/*Δt* curves are at once non-linear and non-superimposing. Therefore, our experimental and numeric results confirm that the LM test is a practical, yet stringent, alternative to detect enzyme inactivation.

**Figure 6.**
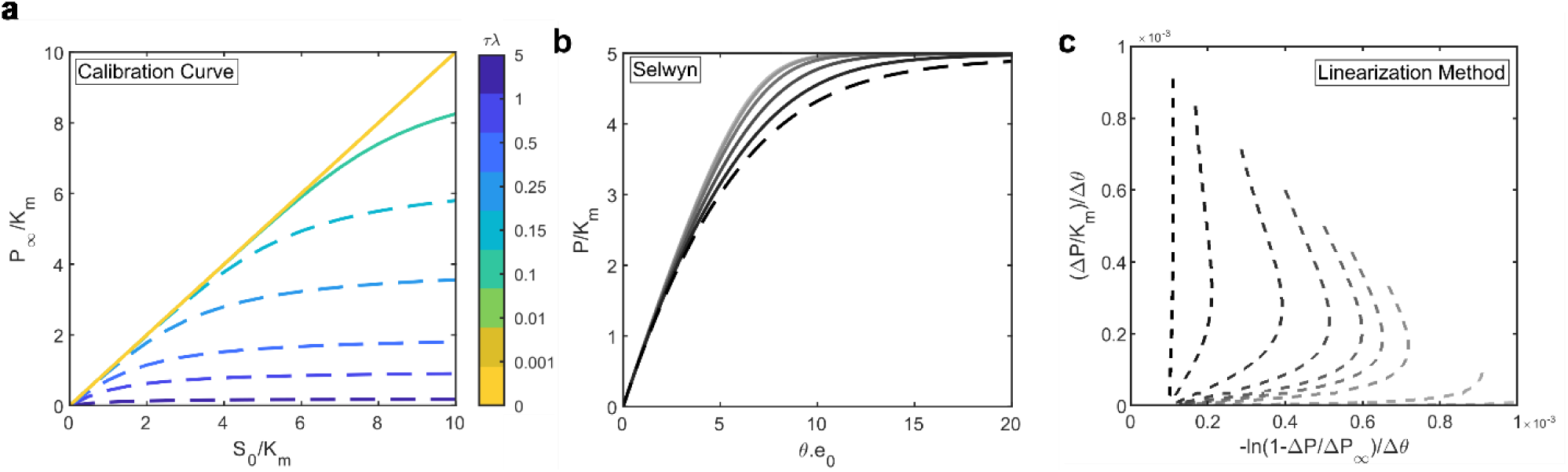
The LM test is highly sensitive to enzyme inactivation. Detection limits of the (a) calibration-curve test, (b) Selwyn test and (c) the LM test according to the theoretical progress curves simulated assuming a first-order decay of enzyme activity (simulation parameters listed in Tables A2–A4). Dashed and solid lines: cases of successful and failed detection, respectively. (a) The calibration-curve test fails to detect inactivation for *τλ* values (color bar) below 0.1, as the measurable and expected values of final product concentration (*P*_∞_ and *S*_0_) start to be undistinguishable. (b) Theoretical progress curves simulated for varying *e*_0_ values for a range of *τλ* values between 10^−3^ and 0.1 (from lighter to darker shades of gray), and *S*_0_/*K*_*m*_ = 5. The Selwyn test fails to detect inactivation for *τλ* < 0.1, as the Selwyn curves are not easily distinguishable. (c) Theoretical LM curves simulated for a reference value of *τλ* = 0.1 and a range of *S*_0_/*K*_*m*_ values between (from lighter to darker shades of gray) 0.1 and 10. This test detects inactivation under conditions of *τλ* = 0.1 and *S*_0_/*K*_*m*_ < 5 for which the calibration-curve test and the Selwyn test have poor sensitivity.

### Unspecific interferences detected in the caspase-3 assay

Faster catalytic reactions are expectable for purified caspase-3 relatively to procaspase-3, which, besides being less active, is present in low concentration in cell extracts. Consequently, the inactivation issues considered for the proenzyme are less important for the purified enzyme - note that the duration of the enzymatic reactions decreases from > 1 day (Figure 3a) to < 1 h (Figure 7a). New challenges for the accurate determination of kinetic parameters are, however, posed by the shorter reaction timescales. This is illustrated in Figure 7a, where the phase of constant velocity is not clearly defined during a stabilization period of ∼10 min. Short periods of normalization of reaction conditions are hardly avoidable even when, as in the present case, the component solutions are pre-equilibrated to the reaction temperature, or when miniaturized high-throughput devices are employed [40]. Small temperature variations markedly influence enzymatic reaction rates and in a *S*_0_-dependent manner [41]. After confirming that the caspase-3 assay is not affected by significant enzyme inactivation (Figure 7b) the reaction phases that succeed the first ∼10 min interval can be analyzed in detail. Because the initial slopes (and the corresponding values of *v*_0_) are probably affected by drifts in the reaction properties, instantaneous rates (*v*_*i*_) obtained upon condition stabilization may be used in MM plots as an approximation to the real value of *v*_0_ (Figure 7c). For rapidly progressing reactions, this procedure raises the doubt of whether the instantaneous substrate concentration is too depleted relative to the initial value (*S*_0_) [42]. Also, it is not granted that the properties of the reaction mixture are completely stabilized during the period of time over which *v*_*i*_ is determined.

**Figure 7.**
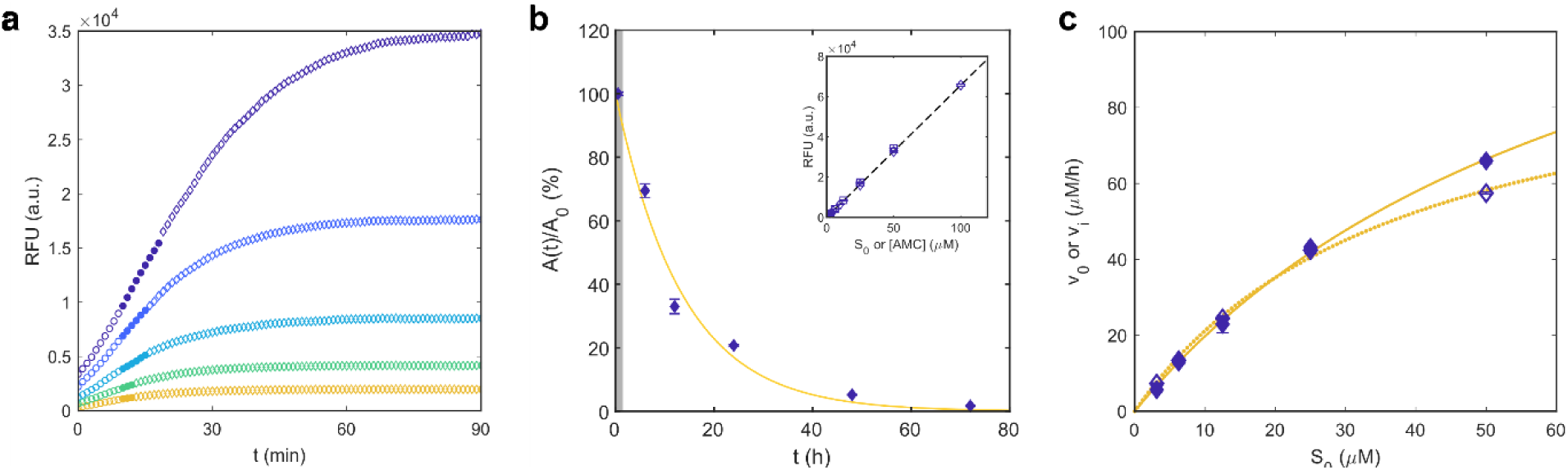
Assay interferences other than enzyme inactivation may affect the initial-rate measurements in the reactions catalyzed by purified caspase-3. (a) Fluorescence increase during the hydrolysis of Ac-DEVD-AMC by 1.0 U of caspase-3 for *S*_0_ values of (from top to bottom) 50, 25, 12.5, 6.125, and 3.125 μM. Circles: data selected for the determination of initial (open symbols) and instantaneous (closed symbols) slopes. (b) Symbols and error bars: means and standard deviations of direct measurements of caspase-3 activity after different periods of incubation in the reaction environment. No significant inactivation occurs within the full reaction timescale (grey area) considered in (a). Solid line: Numerical fit to an exponential decay function (*λ* = 0.074 ± 0.009 h^-1^). Inset: the calibration-curve test confirms the absence of significant enzyme inactivation: the obtained end-point signals (circles) overlie the calibration curve (dashed line) built with fluorescence measurements of standard AMC solutions (squares). (c) Plot of the initial (*v*_0_) and instantaneous (*v*_*i*_) reaction rates as a function of the initial substrate concentration (*S*_0_). The experimental values of *v*_0_ (open symbols) and *v*_*i*_ (closed symbols) are calculated using initial and instantaneous slopes, respectively, as represented in (a). Lines: fit of the experimental data to the MM-like equation. Since both *v*_0_ and *v*_*i*_ are imperfect estimations of the initial rate value (see text for details) the fitted values of *K*_*m*_ = 38.7 ± 6.4 μM and *v*_*max*_ = 103 ± 10 μM/h (dotted line, open symbols) and *K*_*m*_ = 71.9 ± 13.0 μM and *v*_*max*_ = 162 ± 20 μM/h (solid line, closed symbols) are merely indicative.

A better perception of the main experimental outliers can be obtained by representing the progress curves in the modified scale proposed by the LM test (Figure 8a). For each *S*_0_ condition, the initial measurements stand out as evidently separated from the negatively-sloped trend exhibited by most of the subsequent data points. This suggests that the values of *v*_0_ determined from Figure 7a (open circles) and the resulting kinetic analysis in Figure 7c (dotted line) are affected by assay interferences. The fact that no straight line common to all *S*_0_ conditions is clearly defined by the late data points does not have a particular meaning because early errors can propagate throughout the Δ*P*/Δ*t* vs. −ln(1 − *ΔP*/*ΔP*_∞_)/*Δt* curve. Moreover, the final amplification of the instrumental noise is expectable in result of the use of the logarithm in the horizontal axis. Setting the new initial condition to *t*_*i*_ = 10 min not only eliminates the initial outliers but also improves the quality of the linearized plots (Figure 8b). The tendency of the different experimental curves to superimpose in a single straight line further confirms that the activity of caspase-3 remains practically unchanged during the time-course of the reactions. Overall, the first (Figure 8a) and second (Figure 8b) LM representations validate the enzymatic assay for purified caspase-3, although the initial ∼10 min stabilization period should not be considered for analysis. In the determination of instantaneous reaction rates, the used *P*(*t*) data (closed circles in Figure 7a) a1ready integrate the negatively-sloped trends in Figure 8a. The kinetic laws based on *v*_*i*_ measurements (Figure 7c, solid line) should, however, take into to account the depletion of substrate until the moment when the rate is determined [31]. This correction to the MM plots involves replacing the initial substrate concentration *S*_0_ by the instantaneous substrate concentration *S*_*i*_, here estimated using median concentration values for the time interval used for *v*_*i*_ determination. The MM parameters fitted to the *v*_*i*_ vs. *S*_*i*_ data (Figure 8c; *K*_*m*_ = 21.5 μM and *v*_*max*_ = 109 μM/h) are considered valid and free from major assay interferences. In accordance to what is expected for steady-state conditions, the apparent kinetic constants 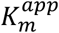 and 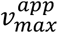 of the LM equation can be approximated by the true parameters *K*_*m*_ and *v*_*max*_. Illustrating this, the experimental LM curves (symbols in Figure 8b) are well described by the theoretical LM curve computed for 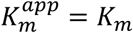 and 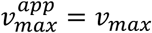 (dashed line in Figure 8b).

**Figure 8.**
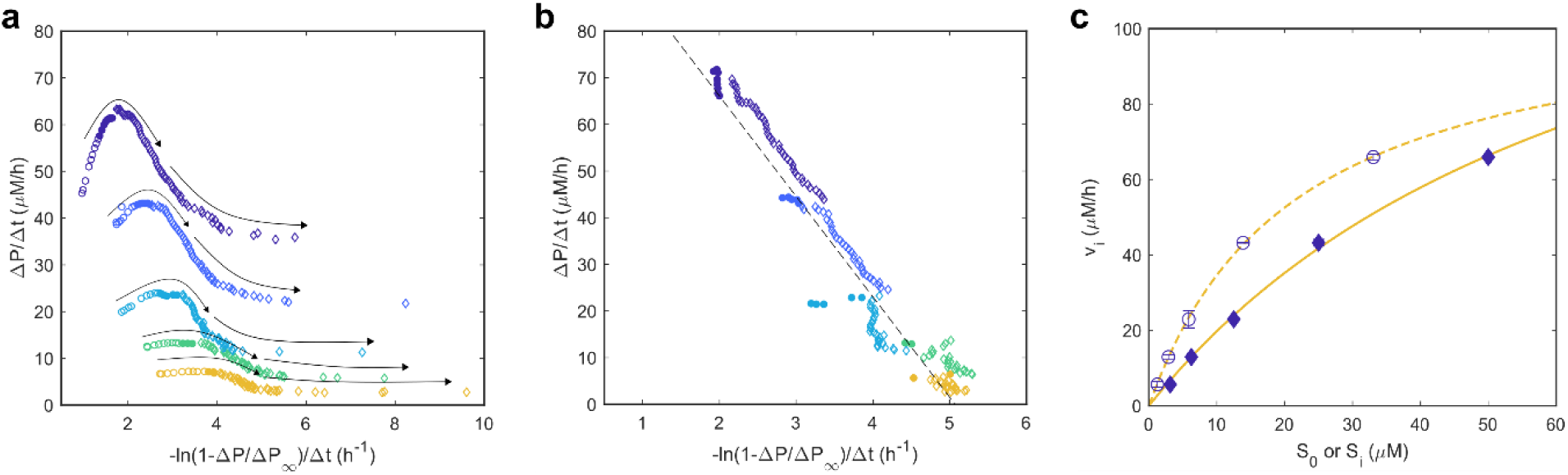
Time-wise variations in solution properties are detected by the LM test. (a) Symbols: progress curves of Ac-DEVD-AMC catalysis by 1.0 U caspase-3 represented in the linearized scale using *t*_*i*_ = 0 min (color-coded as in Figure 7a). Arrows: visual reference indicating the reaction time-course. The initial experimental outliers (large open symbols) show up detached from the negatively-sloped trends. The final scattering of the data results from the amplification of random errors and instrumental noise. (b) Symbols: linearized progress curves obtained after discarding the initial outlier points (*t*_*i*_ = 10 min). For clarity, only data corresponding to 95% reaction completion are represented. Dashed line: representation of Eq. 2 after replacing 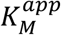 and 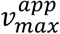 by the values of *K*_*m*_ and *v*_*max*_ determined by independent methods in (c). (c) Symbols and error bars: means and standard deviations of the instantaneous reaction rates (*v*_*i*_) represented as a function of *S*_0_ (closed symbols) and *S*_*i*_ (open symbols). Lines: numerical adjustment of the MM-like equation. The *v*_*i*_ vs. *S*_0_ data and numerical fit (closed symbols and solid line) are the same as in Figure 7c (shown here as visual reference). Dashed line: numerical fit of the *v*_*i*_ vs. *S*_*i*_ data; the fitted results (*K*_*m*_ = 21.5 ± 0.9 μM and *v*_*max*_ = 109 ± 2 μM/h) are not affected by major assay interferences as they successfully describe the experimental LM curves in (b).

### Inhibition of α-thrombin

The presence of unaccounted enzyme modifiers in the assay solution is another possible interference associated to the use of crude enzyme preparations and cell extracts. The detection of enzyme modulation effects by the LM test is here demonstrated for the inhibition of the amydolytic activity of human α-thrombin by a synthetic variant of an anticoagulant produced by *D. andersoni*. Since the inhibitory effect of nanomolar concentrations of the enzyme-modifier is known beforehand (Figure A1, Appendix), the LM test is applied to identify the fingerprints left by enzyme modifiers and to illustrate how cautiously initial-rate measurements should be used during the characterization of inhibition mechanisms.

If (i) quasi-equilibrium conditions are rapidly attained and (ii) the concentration of inhibitor is significantly higher than the concentration of enzyme, progress curves measured in the presence of competitive, uncompetitive or mixed inhibition are still numerically described by Eq. 2, with 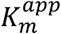 and 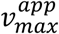 being affected by the concentration of inhibitor(s). Consequently, the LM analysis may fail to detect linear inhibition effects provoked by solution contaminants. On the positive side, quasi-equilibrium linear inhibition is a particular case of the general modifier mechanism [43], whose rate equations frequently contain squared concentration terms recognizable as deviations from the LM equation. As such, it is conceivable that the new method can also be used for identifying non-standard inhibition mechanisms and not only for assay validation purposes. The complexity of this subject greatly increases as other LM-detectable mechanisms of product inhibition, slow-onset inhibition, substrate competition, allosterism, etc., are considered. Presently, we apply the LM test as a “black box” test to α-thrombin-catalyzed reactions inhibited by a synthetic variant of an anticoagulant produced by *D. andersoni*, and we leave the fundamental characterization of the inhibition mechanism to future research. This model system is useful to illustrate how the presence in vestigial amounts (0.40 nM) of a given compound might be revealed by characteristic kinetic signatures left in LM curves. The sigmoid-shaped onset of the α-thrombin progress curves (Figure 9a) is admissible under conditions of *S*_0_ ≤ *E*_0_ that do not apply here [31]. Once again, the ill-defined initial rates can admittedly result from the gradual stabilization of the experimental conditions and not necessarily from the presence of enzyme modifiers. Yet, unlike what was observed for capase-3, the first LM representation (Figure 9b) suggests that stabilization periods much longer than 10 min are required for the emergence of negatively-sloped LM curves. Even admitting a stabilization period of 30 min in the second LM representation (Figure 9c), no superimposable trend is clearly defined by the individual curves obtained at the different *S*_0_ conditions. This means that, in the case of α-thrombin, imperfect temperature control during the initial reaction phases cannot be used to explain the inconsistent LM curves obtained afterwards. Taken together, the long initial periods evidencing positive LM slopes and the persisting lack of a well-defined trend common to the different *S*_0_ conditions indicate possible deviations from the basic Briggs-Haldane mechanism. After dismissing the hypothesis of enzyme inactivation (which can be done using the calibration curve test), the presence of high-affinity enzyme modifiers is a strong possibility to be considered even when their action is through a direct inhibition mechanism. In fact, rate equations containing squared concentration terms are not exclusive of hyperbolic modifiers but are also expected for tight-binding linear inhibitors occurring at concentrations in the order of magnitude of *E*_0_ or lower [44–46]. For this reason, unsuspected contaminants that are also tight-binding modifiers will have their effects uncovered by representing the measured progress curves in LM coordinates. Although the saturation plots obtained using initial (index 0) or instantaneous (index *i*) variables correspond to typical MM curves (Figure 9d), the fitted parameters are, in both cases, of ambiguous physical meaning. The *v*_0_ vs. *S*_0_ analysis (solid lines in Figure 9b and Figure 9c) is clearly affected by assay interferences as the theoretical LM curve (solid line in Figure 9c) fails to intercept the experimental LM curves. When instantaneous measurables are analyzed (dashed lines in Figure 9c and Figure 9d), the theoretical LM curve is able to intercept, at least in part, the experimental results obtained for each *S*_0_ condition (Figure 9c); even so, the α-thrombin enzymatic assay is considered non-compliant with the LM prerequisites because the individual LM time-course trends (indicated by symbol size increase in Figure 9c) are divergent from the overall straight line suggested by the theoretical LM curve (dashed line in Figure 9c). This didactic example serves to demonstrate that assay interferences cannot be diagnosed solely based on the quality of numerical adjustments to the MM equation. In contrast, kinetic effects caused by very small amounts of either linear or hyperbolic inhibitors can be detected by the LM test.

**Figure 9.**
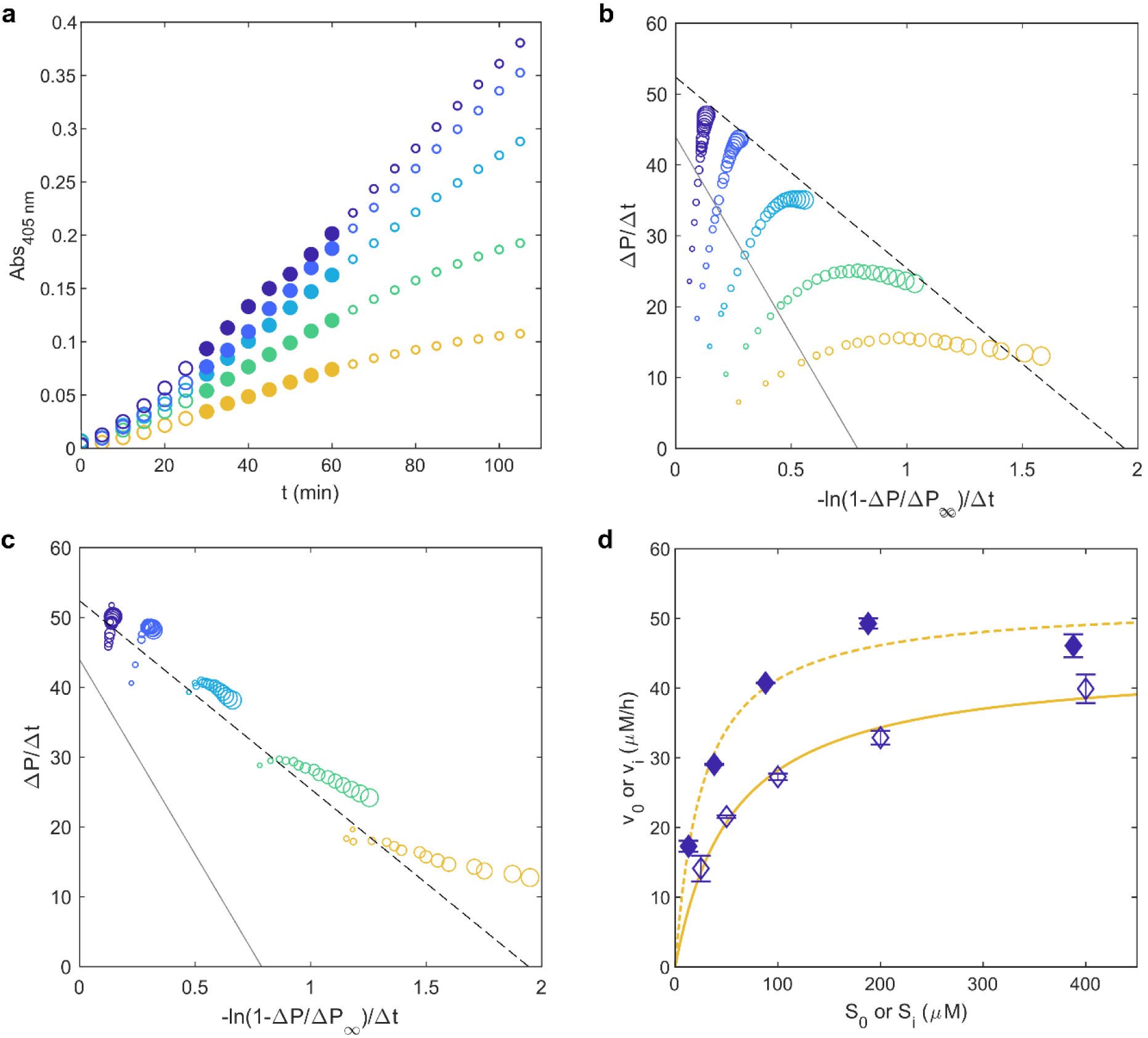
The LM test detects timewise effects induced by enzyme modifiers. Absorbance (*Abs*_405*nm*_) increase during the catalysis of Tos-Gly-Pro-Arg-pNA by 0.15 nM α-thrombin in the presence of 0.40 nM inhibitor for substrate concentration of (from top to bottom) 400, 200, 100, 50 and 25 μM. Large symbols: data selected for the determination of initial (large open symbols) and instantaneous (closed symbols) slopes. (b) The same progress curves are represented in the linearized scale using *t*_*i*_ = 0 min. (c) Symbols: LM progress curves obtained after discarding the initial outlier points (*t*_*i*_ = 30 min). (b,c) Lines: representation of Eq. 2 after replacing 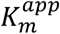 and 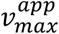 by the values of *K*_*m*_ and *v*_*max*_ determined in (d) using initial (solid line) and instantaneous (dashed line) measurables. Symbol size increases with the time-course of the reaction. (d) Symbols and error bars: means and standard deviations of the initial reaction rates (*v*_0_) and instantaneous reaction rates (*v*_*i*_) represented as a function of *S*_0_ (open symbols) and *S*_*i*_ (closed symbols), respectively. Lines: using the MM-like equation to fit *v*_0_ vs. *S*_0_ data (solid line, *K*_*m*_ = 55.8 ± 9.3 μM and *v*_*max*_ = 44.1 ± 2.3 μM/h) and *v*_*i*_ vs. *S*_*i*_ data (dashed line, *K*_*m*_ = 26.9 ± 7.2 μM and *v*_*max*_ = 52.6 ± 3.4 μM/h). The LM test is not passed because the individual trends in (c) fail to converge into a single overall straight line.

### The LM test as a routine quality check

The LM test is suitable for routine use in enzymatic assay validation and to decide which optimization steps should be taken in order to improve reproducibility and accuracy. As summarized in Figure 10, its application is based on the LM representations of reaction progress curves obtained for a fixed value of *E*_0_ and varying *S*_0_. First, the coordinates Δ*P*/Δ*t* and −ln(1 − *ΔP*/*ΔP*_∞_)/*Δt* are computed using the beginning of the measurements (*t*_*i*_ = 0 min) as initial condition for identifying the initial period of stabilization of reaction conditions. If the obtained LM curves are negatively-sloped straight lines and tend to superimpose since the beginning of the reaction, the test is passed. If evident deviations from the ideal trend are found to occur, new coordinates must be computed using as initial reference any instant subsequent to the period of stabilization (*t*_*i*_ > 0 min). If the LM test is still not passed, the presence of enzyme inactivation can be tested using the calibration-curve test. Some assay interferences can be minimized by increasing the concentration and/or purity of the enzyme; the presence of unspecific assay interferences must also be checked here. This process can be repeated iteratively until the enzymatic assay is fully optimized. Next, we present additional practical guidelines to be considered during the systematization of the new method.

**Figure 10.**
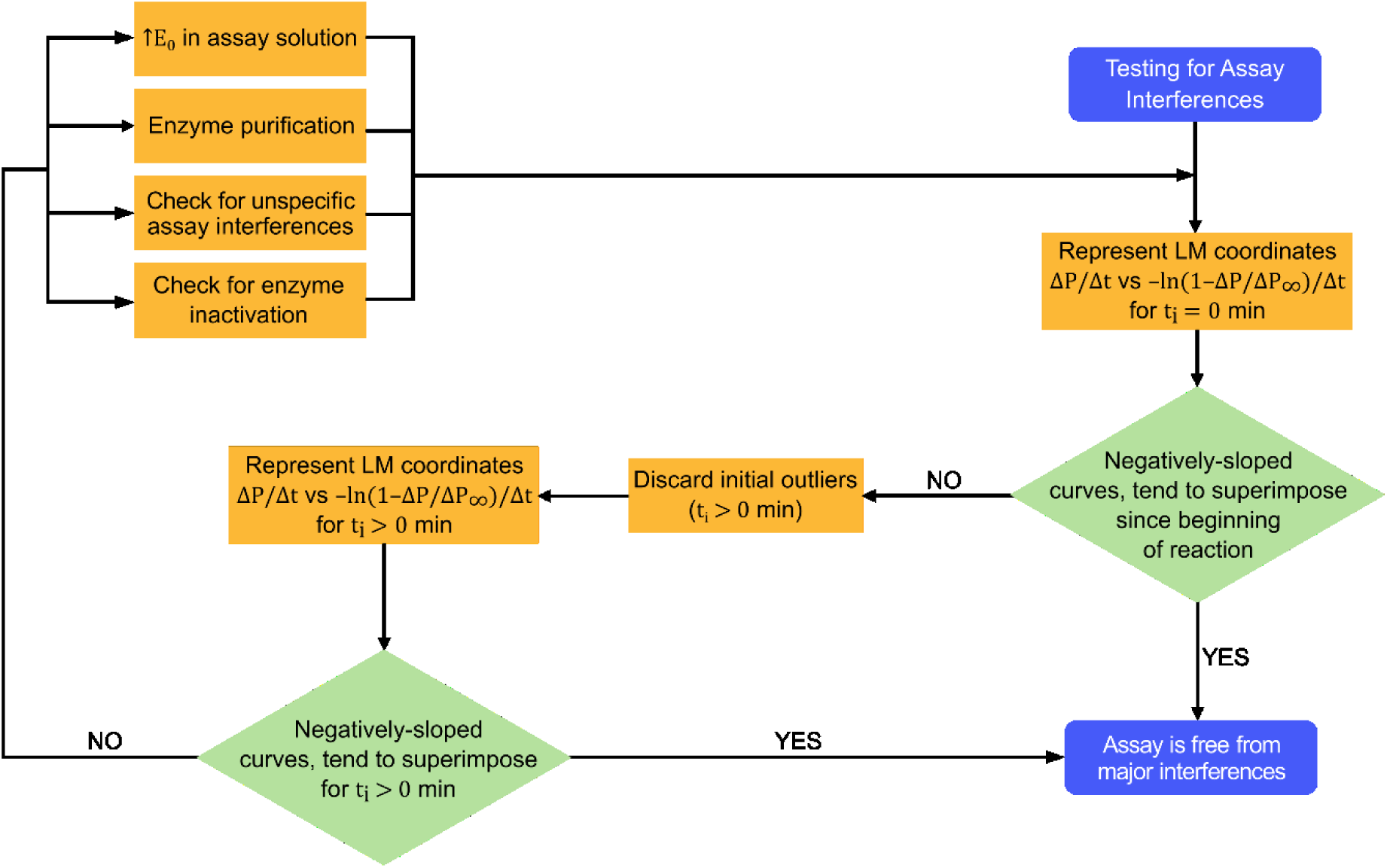
Flowchart depicting the main steps of the LM test for the identification of assay interferences.

More complete information is provided by the LM test when full progress curves are measured. Note, for example, that the *ΔP*/*Δt* coordinate does not change during the whole initial period of constant velocity. If, in a limit case, all points are collected within constant velocity timeframes, no individual trends will be defined for each substrate concentration and the result of the test will be solely dictated by the linearity of the overall trend. Time-dependent changes in enzymatic activity can thus be missed when only a part of the reactions is analyzed. Letting the reaction proceed until its end also allows identifying the differences between *P*_∞_ and *S*_0_ values upon which the calibration-curve test is based. Although less sensitive than the LM test, the calibration-curve test is specific for enzyme inactivation and, for this reason, is indicated for the preliminary assessments of this type of interference. Nonetheless, as shown for the case of α-thrombin, full progress curves are not mandatory for the application of the LM test.

Small errors in reagent handling give rise to differences between the values of *P*_∞_ and *S*_0_ that are only evident at the end of the reaction. Such unspecific interferences are detected by the LM test even if the final product concentrations are not known, and *S*_0_ values have to be used instead of *P*_∞_ to compute the −ln(1 − *ΔP*/*ΔP*_∞_)/*Δt* coordinate. In the numerical example given in the Appendix (Figure A2), a random error of 5% in the value of *S*_0_ generates, since the beginning of the reaction, a clear deviation of the biased LM curve relatively to the overall trend defined by the other LM curves. As the reaction progresses in time, the differences become more pronounced and even the initial linearity is lost. A consequence of the Walker and Schmidt-type linearization adopted by the LM equation [26], this high responsiveness to random experimental errors is helpful for controlling the validity of each reaction rate measurement.

## Conclusions

A new method to identify assay interferences is proposed based on a modified version of the integrated MM equation. To pass the so called “LM test”, progress curves measured at different substrate concentrations and represented in linearized Δ*P*/Δ*t* vs. −ln(1 − *ΔP*/*ΔP*_∞_)/*Δt* coordinates should superimpose in a single, negatively-sloped straight line. The proposal of this new method follows from the recently obtained closed-form solution of the Briggs and Haldane reaction mechanism [31, 32]. Some of the modifications now introduced to the integrated MM equation allow time-course kinetic analysis to be carried out in the range of conditions commonly adopted for steady-state analysis (*E*_0_ ≪ *S*_0_ + *K*_*M*_). The illustrative examples of enzymatic reactions catalyzed by procaspase-3, capase-3 and α-thrombin highlight different aspects that can stealthily influence the quality of enzymology data. Initial rate measurements during the catalysis of Ac-DEVD-AMC by procaspase-3 in yeast cell extracts are strongly affected by progressive enzyme inactivation, promptly detected by the LM test independently of whether such suspicion exists *a priori* or not. The Selwyn test, a reference method to identify enzyme inactivation [28, 47], is less sensitive than the LM test and requires additional experiments whenever enzyme inactivation is somehow suspected to occur. The catalysis of Ac-DEVD-AMC by purified caspase-3 was used to demonstrate that non-conformities in the first LM representation are a possible indication that the experimental conditions were not yet stabilized at the beginning of the measurements. In fact, the LM representation obtained after discarding the initial stabilization period validated the caspase-3 assay, which was then used to determine unbiased MM parameters based on instantaneous values of substrate concentration and reaction rate. Finally, the kinetic fingerprints left by enzyme modifiers were revealed during the inhibition of α-thrombin by nanomolar concentrations of a synthetic anticoagulant. This example illustrated that high-affinity contaminants may affect enzyme kinetics in a hard to detect way unless LM plots are represented and individual LM curves are compared with the overall trend.

Because stringent criteria are adopted, nonstandard catalytic reactions (characterized by multiple active sites, multiple substrates, product inhibition, hyperbolic inhibition, tight-binding inhibition, enzyme inactivation, etc.), or that are influenced by instrumental noise and poor automatic control will probably not pass the LM test. This fine quality control is crucial for the success of quantitative kinetic analysis since important mechanistic nuances “may play a subordinated role with respect to even modest mistakes in reagent handling, (…) instrumental noise and others” [48]. In line with current initiatives to improve enzymology data reporting [49, 50], this new method is expected to contribute in improving the reproducibility of kinetic data, thus increasing the impact of fundamental and applied research in fields such as enzyme engineering, systems biology and drug discovery.

## Acknowledgements

This work was financed by (i) FEDER—Fundo Europeu de Desenvolvimento Regional funds through the COMPETE 2020—Operacional Programme for Competitiveness and Internationalisation (POCI), Portugal 2020, and by Portuguese funds through FCT—Fundação para a Ciência e a Tecnologia/Ministério da Ciência, Tecnologia e Ensino Superior in the framework of the projects POCI-01-0145-FEDER-031173 (PTDC/BIA-BFS/31173/2017), POCI-01-0145-FEDER-031323 (“Institute for Research and Innovation in Health Sciences”) and POCI-01-0145-FEDER-006939 (Laboratory for Process Engineering, Environment, Biotechnology and Energy – UID/EQU/00511/2013), POCI-01-0145-FEDER-007728, (ii) FEDER through Norte Portugal Regional Operational Programme (NORTE 2020), under the PORTUGAL 2020 Partnership Agreement in the framework of Project Norte-01-0145-FEDER-000008 and by national funds (PIDDAC) through FCT project “LEPABE-2-ECO-INNOVATION” – NORTE-01-0145-FEDER-000005, and by (iii) National Funds (FCT/MEC), under the Partnership Agreement PT2020 UID/MULTI/04378/2013 and the projects (3599-PPCDT) PTDC/DTP-FTO/1981/2014 – POCI-01-0145-FEDER-016581.

PhD fellowships SFRH/BD/109324/2015 (MFP) and SFRH/BD/119144/2016 (HR), and Postdoctoral fellowship SFRH/BPD/108004/2015 (J.R.-R.) from FCT – Fundação para a Ciência e a Tecnologia (Programa Operacional Capital Humano (POCH), UE) are acknowledged. Financial support was provided through the Doctoral Program in Biomedical Sciences (ICBAS-UP) and BiotechHealth Programme (ICBAS-UP/FFUP).

## Appendix

### A1. Irreversible enzyme inactivation - numeric solutions

The Briggs-Haldane reaction scheme comprises the binding of substrate (*S*) to free enzyme (*E*) to give rise to the enzyme-substrate complex (*ES*), from which the catalytic product (*P*) is formed and released, thereby regenerating the free enzyme. Irreversible enzyme inactivation can be accounted for in this scheme by considering that free and bound enzyme decay irreversibly into the inactive forms (*E*\* and *ES*\*). For simplicity we will assume that both processes are well described by the same first-order rate constant (*λ*):

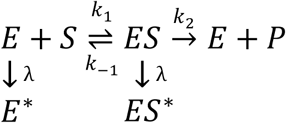

This reaction scheme is mathematically described by the following system of first-order differential equations:

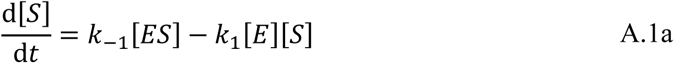

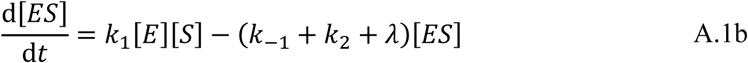

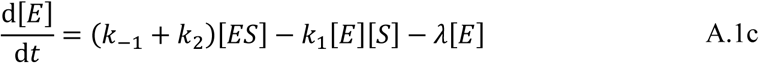

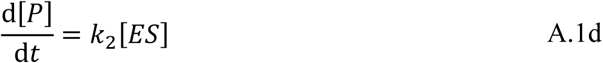

subject to the initial conditions ([*S*], [*ES*], [*E*], [*P*]) = (*S*_0_, 0, *E*_0_, 0) and to the mass conservation laws *E*_0_ = [*E*] + [*ES*] + [*E*\*] + [*ES*\*] and *S*_0_ = [*S*] + [*ES*] + [*P*]. The concentrations of the different species can be normalized by the Michaelis constant *K*_*m*_ = (*K*_-1_+*K* _2_)/ *K* _1_ as *s* = [*S*]/*K*_*m*_, *c* = [*ES*]/*K*_*m*_, *e* = [*E*]/*K*_*m*_ and *p* = [*P*]/*K*_*m*_, and expressed as a function of the modified timescale *θ* = *K* _2_*t*:

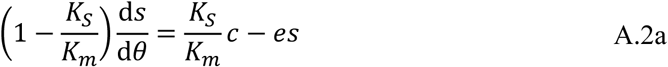

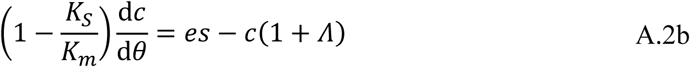

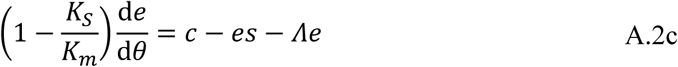

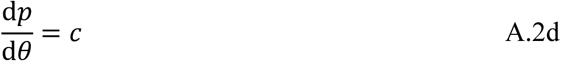

where *K*_*S*_ = *K*_-1_/ *K* _1_ is the dissociation constant of the enzyme-substrate complex, and *Λ* is a normalization of *λ*:

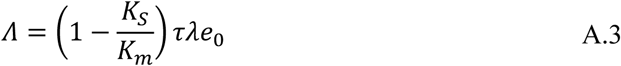

The system of ordinary differential equations comprising Eqs. A.2a-A.2d was numerically solved using the ode15s solver of Mathworks® MATLAB 2018a (Natick, MA, USA) for the sets of simulation parameters summarized in Tables A1, A3, and A4.

### A2. Irreversible enzyme inactivation -approximate analytical solution

The instantaneous concentration of total active enzyme (*E*_*a*_) is given by the sum of active enzyme in its free (*E*) and bound state (*ES*). Since both forms are assumed to inactivate at the same rate, the evolution of *E*_*a*_ over time assumes the form of an exponential decay function:

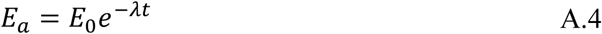

For sufficiently high values of initial substrate concentration *S*_0_, Pinto, et al. [31] showed that the reaction rate equation (Eq. S.1d) can be rewritten as:

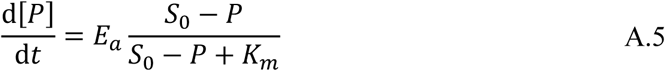

After replacing the time-dependent definition of *E*_*a*_ (Eq. A.4), the analytical integration of A.5 gives the evolution of [*P*] as a function of the time *t* in the presence of enzyme inactivation:

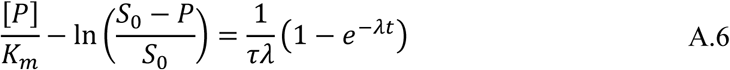

with *τ* = *K*_*m*_/(*K*_2_ *E*_0_). The limit of Eq. A.6 for *t* → +∞ defines the final concentration of obtained product *P*_∞_ as a function of *S*_0_:

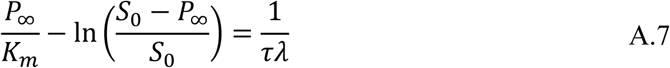

The Lambert ω function is a built-in function in mathematical software that satisfies the transcendental equation *ω*(*x*)*e*^*ω*(*x*)^ = *x*. It is used to express Eqs. A.6 and A.7 as closed-form solutions, solving for [*P*] (Eq. A.8) and *P*_∞_ (Eq. A.9):

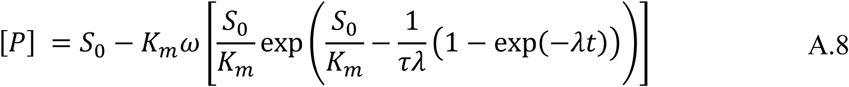

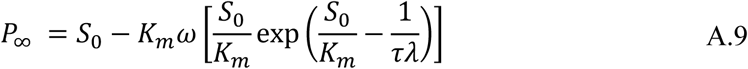

Equation A.9 was used to simulate *P*_*∞*_/*K*_*m*_ vs *S*_0_/*K*_*m*_ curves for the sets of simulation parameters summarized in Table A2.

### A3. Reaction rate analysis

Initial reaction rates (*v*_0_) of progress curves for varying substrate concentrations were obtained and analyzed to determine kinetic parameters *K*_*m*_ and *v*_*max*_ = *K*_2_*E*_0_. Data were fitted to the MM equation (Eq. A.10) by the non-linear least-squares method.

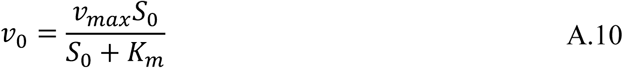

For procaspase-3 data, initial time intervals with approximate linear behavior were chosen for *v*_0_ determination (Table A5). For recombinant caspase-3 and α-thrombin data, initial velocity and instantaneous velocity values (*v*_0_ and *v*_*i*_, respectively) were determined. The selected time intervals used in each case are listed in Table A5. Instantaneous substrate concentrations *S*_*i*_ were determined for the median points of the intervals chosen for *v*_*i*_ determination. Non-linear least-squares fitting of *v*_*i*_ vs *S*_*i*_ was performed using the MM-like equation described by Pinto, et al. [31]:

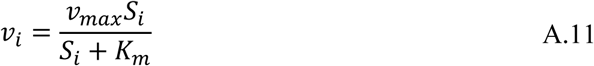

### A4. Figures

**Figure A1.**
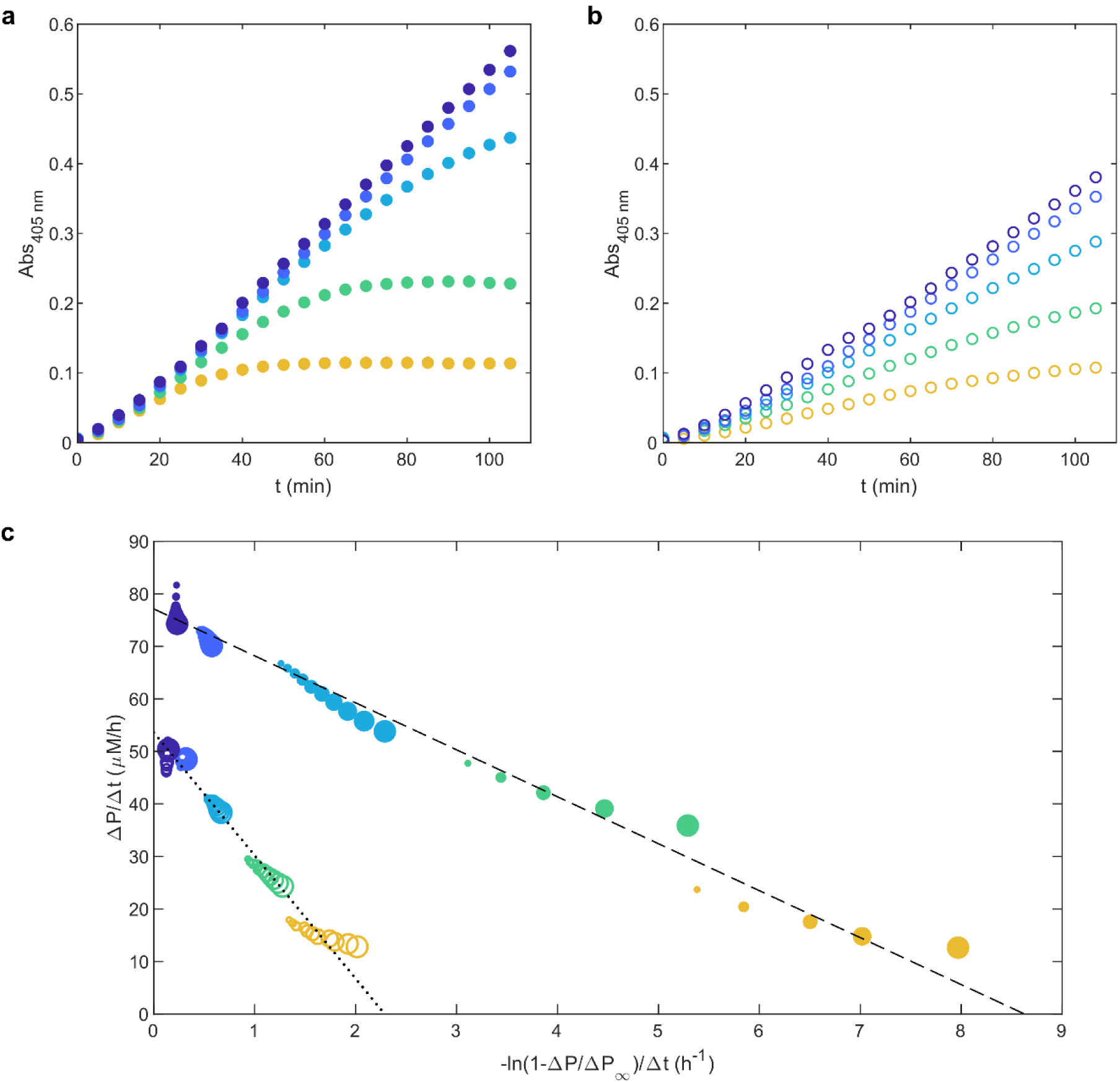
The presence of vestigial amounts (0.40 nM) of a synthetic variant of an anticoagulant produced by *D. andersoni* inhibits the catalysis of Tos-Gly-Pro-Arg-p-NA by 0.15 nM α-thrombin. (a and b) Absorbance (*Abs*_405*nm*_) increase measured in the absence (a) and presence (b) of inhibitor for substrate concentrations of (from top to bottom) 400, 200, 100, 50 and 25 μM. (c) The same progress curves are represented according to the linearization method (LM) scale using *t*_*i*_ = 30 min (color-coded as in Figures A1a and A1b). Symbol size increases according to the time-course of the reaction. Lines: representation of the theoretical LM equation (Eq. 2) using 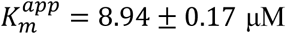 and 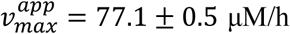 (dashed line) and 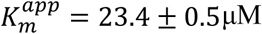 and 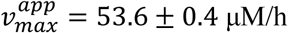 (dotted line).

**Figure A2.**
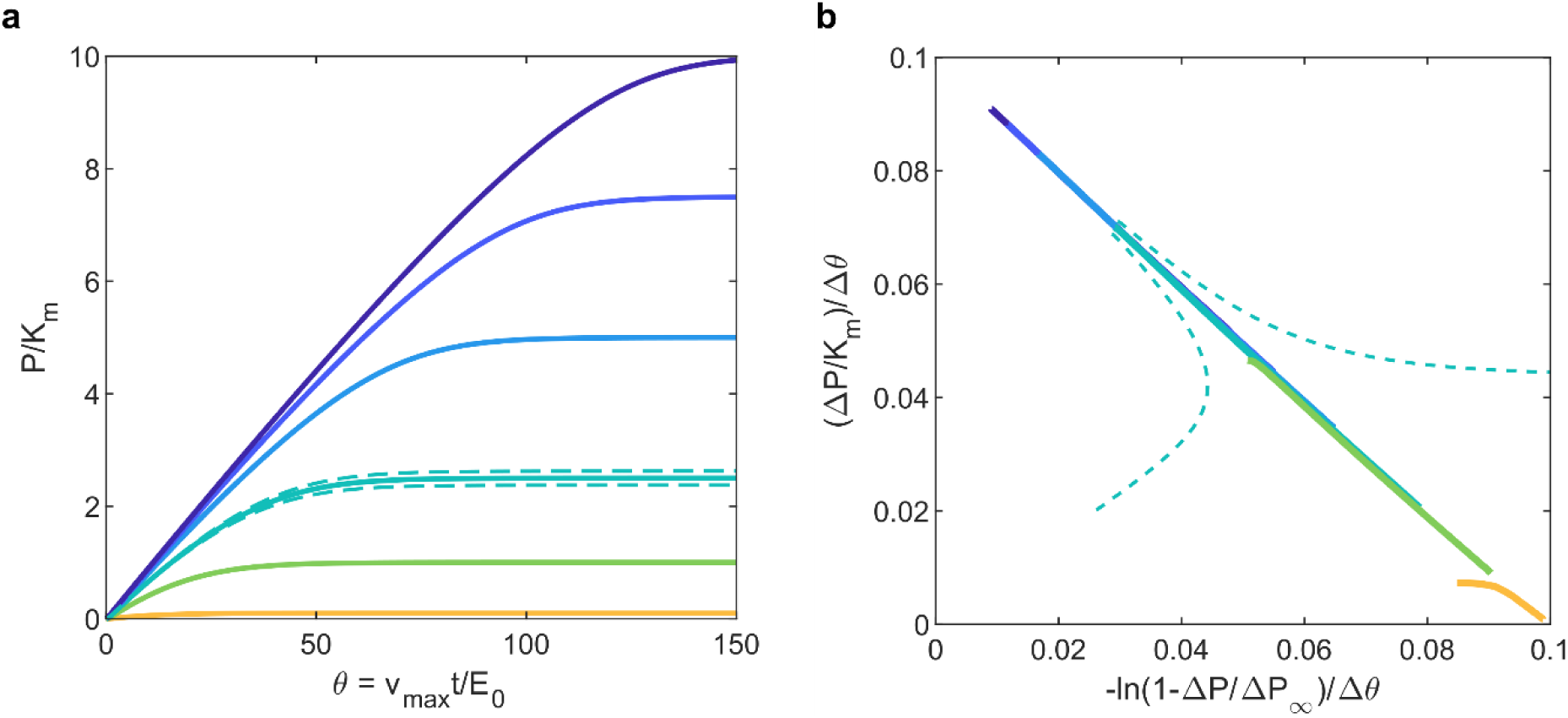
Small errors in reagent handling can be detected by the LM test. (a) Solid lines: theoretical progress curves calculated as described in Figure 2a for *S*_0_/*K*_*m*_ values of (from top to bottom) 10, 7.5, 5, 2.5, 1, 0.1. Dashed lines: theoretical progress curves simulated for a random error of ±5% in the value of *S*_0_/*K*_*m*_ = 2.5. (b) Lines: the same progress curves are represented in the LM scale using *t*_*i*_ = 0, and *P*_∞_ = *S*_0_. A clear deviation of the biased LM curves relatively to the overall trend can be identified since the beginning of the reaction.

### A5. Tables

**Table A1.** Model parameters used for simulation of reaction progress curves and LM curves presented in Figure 2.

| $e_0 = E_0/K_m$ | $S_0/K_m$ | $K_s/K_m$ | $\lambda/k_2$ | $\tau\lambda = (\lambda/k_2)(1/e_0)$ |
| --- | --- | --- | --- | --- |
| 0.1 | 0.1 | 0 | 0 | 0 |
|  | 1 |  |  |  |
|  | 2.5 |  |  |  |
|  | 5 |  |  |  |
|  | 7.5 |  |  |  |
|  | 10 |  |  |  |

**Table A2.** Model parameters used for simulation of *P*_∞_/*K*_*m*_ vs *S*_0_/*K*_*m*_ curves presented in Figure 6a.

| $e_0 = E_0/K_m$ | $S_0/K_m$ | $K_s/K_m$ | $\lambda/k_2$ | $\tau\lambda = (\lambda/k_2)(1/e_0)$ |
| --- | --- | --- | --- | --- |
| $10^{-3}$ | [0,10] | 1 | 0 | 0 |
| | | | $10^{-6}$ | $10^{-3}$ |
| | | | $10^{-5}$ | 0.01 |
| | | | $10^{-4}$ | 0.1<sup>a</sup> |
| | | | $1.5 \times 10^{-4}$ | 0.15 |
| | | | $25 \times 10^{-4}$ | 0.25 |
| | | | $5 \times 10^{-4}$ | 0.5 |
| | | | $1 \times 10^{-3}$ | 1 |
| | | | $5 \times 10^{-3}$ | 5 |

<sup>a</sup>Inactivation effects are poorly detected by the calibration-curve test for reference values of $\tau\lambda < 0.1$ .

**Table A3.**
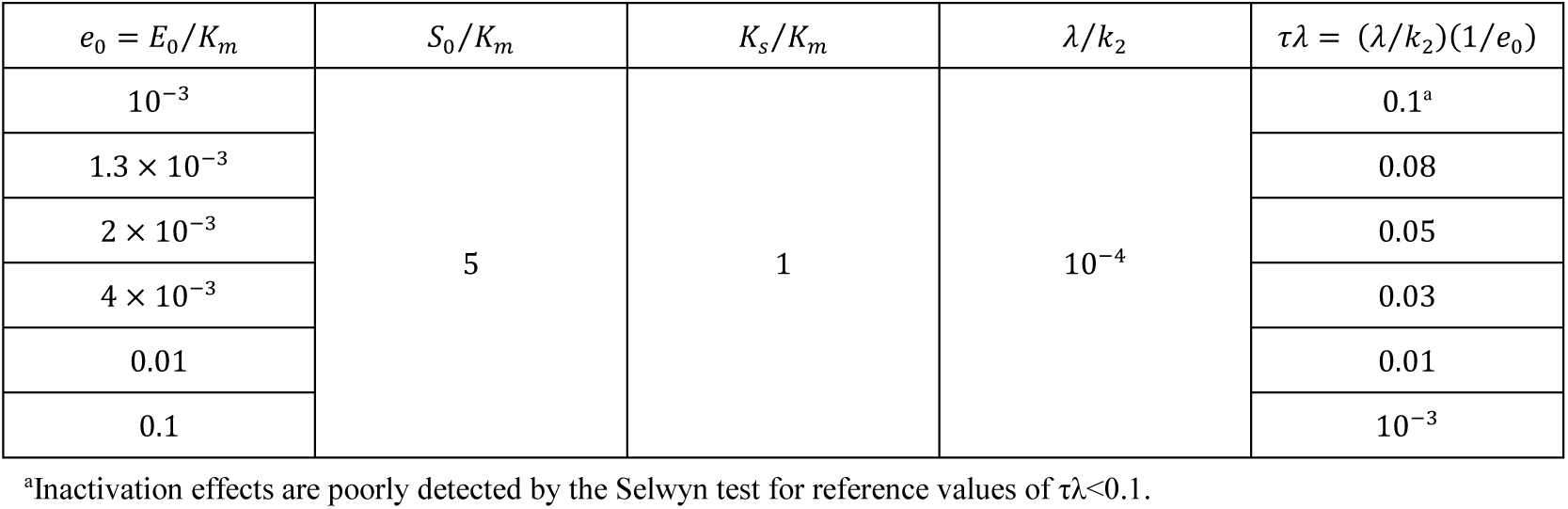
Model parameters used for simulation of reaction progress curves presented in Figure 6b.

| $e_0 = E_0/K_m$ | $S_0/K_m$ | $K_s/K_m$ | $\lambda/k_2$ | $\tau\lambda = (\lambda/k_2)(1/e_0)$ |
| --- | --- | --- | --- | --- |
| $10^{-3}$ | 5 | 1 | $10^{-4}$ | 0.1<sup>a</sup> |
| $1.3 \times 10^{-3}$ | | | | 0.08 |
| $2 \times 10^{-3}$ | | | | 0.05 |
| $4 \times 10^{-3}$ | | | | 0.03 |
| 0.01 |  |  |  | 0.01 |
| 0.1 | | | | $10^{-3}$ |

<sup>a</sup>Inactivation effects are poorly detected by the Selwyn test for reference values of $\tau\lambda < 0.1$ .

**Table A4.**
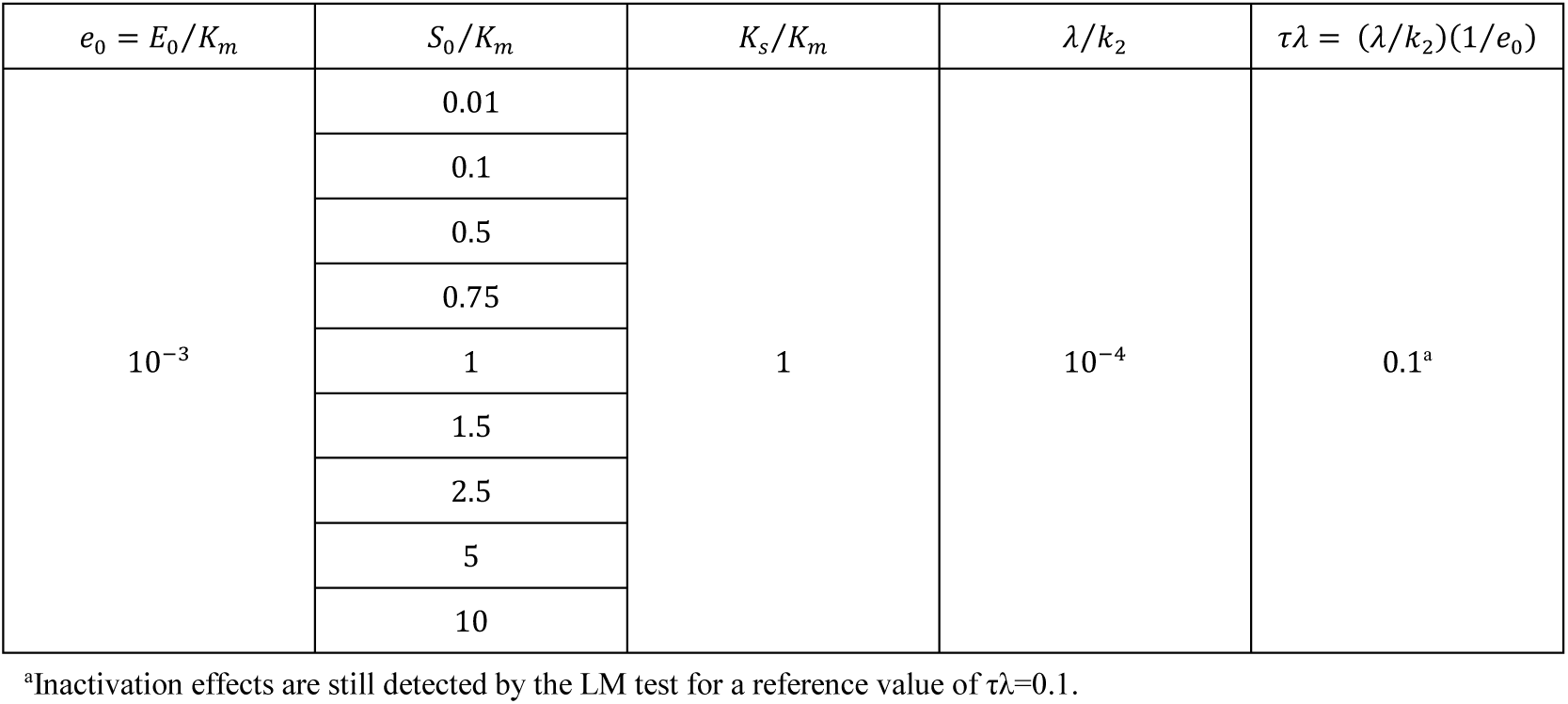
Model parameters used for simulation of LM curves presented in Figure 6c.

| $e_0 = E_0/K_m$ | $S_0/K_m$ | $K_s/K_m$ | $\lambda/k_2$ | $\tau\lambda = (\lambda/k_2)(1/e_0)$ |
| --- | --- | --- | --- | --- |
| $10^{-3}$ | 0.01 | 1 | $10^{-4}$ | 0.1<sup>a</sup> |
|  | 0.1 |  |  |  |
|  | 0.5 |  |  |  |
|  | 0.75 |  |  |  |
|  | 1 |  |  |  |
|  | 1.5 |  |  |  |
|  | 2.5 |  |  |  |
|  | 5 |  |  |  |
|  | 10 |  |  |  |

<sup>a</sup>Inactivation effects are still detected by the LM test for a reference value of $\tau\lambda = 0.1$ .

**Table A5.** Time intervals employed for determination of initial reaction velocity *v*_0_ and instantaneous reaction velocity *v*_*i*_ by linear regression for procaspase-3, recombinant caspase-3 and α-thrombin enzymatic assays.

|  |  |  |  |  |  |  |  |  |  |  |
| --- | --- | --- | --- | --- | --- | --- | --- | --- | --- | --- |
| Procaspase-3 | $S_0$ ($\mu\text{M}$) | 3.125 | 6.25 | 12.5 | 25 | 50 | 100 | 150 | 200 | 300 |
| | $t_{int}$ (h), $v_0$ | [0,0.05] | [0,0.05] | [0,0.07] | [0,0.125] | [0,0.25] | [0,5] | [0,75] | [0,1] | [0,2] |
| Recombinant caspase-3 | $S_0$ ($\mu\text{M}$) | 3.125 | | 6.25 | | 12.5 | | 25 | | 50 |
| | $t_{int}$ (h), $v_0$ | [0,0.15] | | [0,0.15] | | [0,0.15] | | [0,0.15] | | [0,0.15] |
| | $t_{int}$ (h), $v_i$ | [0.15,0.20] | | [0.15,0.20] | | [0.15,0.25] | | [0.15,0.25] | | [0.15,0.30] |
| α-thrombin<br>$I = 0.40$ nM | $S_0$ ($\mu\text{M}$) | 25 | | 50 | | 100 | | 200 | | 400 |
| | $t_{int}$ (h), $v_0$ | [0,30] | | [0,30] | | [0,30] | | [0,30] | | [0,30] |
| | $t_{int}$ (h), $v_i$ | [30,40] | | [30,45] | | [30,60] | | [30,60] | | [30,60] |

